# From drift to draft: How much do beneficial mutations actually contribute to predictions of Ohta’s slightly deleterious model of molecular evolution?

**DOI:** 10.1101/681866

**Authors:** Jun Chen, Sylvain Glémin, Martin Lascoux

## Abstract

Since its inception in 1973 the slightly deleterious model of molecular evolution, aka the Nearly Neutral Theory of molecular evolution, remains a central model to explain the main patterns of DNA polymorphism in natural populations. This is not to say that the quantitative fit to data is perfect. In a recent study Castellano *et al.* (2018) used polymorphism data from *D. melanogaster* to test whether, as predicted by the Nearly Neutral Theory, the proportion of effectively neutral mutations depends on the effective population size (N_e_). They showed that a nearly neutral model simply scaling with N_e_ variation across the genome could not explain alone the data but that consideration of linked positive selection improves the fit between observations and predictions. In the present article we extended their work in two main directions. First, we confirmed the observed pattern on a set of 59 species, including high quality genomic data from 11 animal and plant species with different mating systems and effective population sizes, hence *a priori* different levels of linked selection. Second, for the 11 species with high quality genomic data we also estimated the full Distribution of Fitness Effects (DFE) of mutations, and not solely the DFE of deleterious mutations. Both N_e_ and beneficial mutations contributed to the relationship between the proportion of effectively neutral mutations and local N_e_ across the genome. In conclusion, the predictions of the slightly deleterious model of molecular evolution hold well for species with small N_e_. But for species with large N_e_ the fit is improved by incorporating linked positive selection to the model.

## Introduction

The year 2018 saw the celebration of the 50^th^ anniversary of the Neutral Theory of molecular evolution (called simply the Neutral Theory thereafter). At 50 years of age, the Neutral Theory is still shrouded in controversies, some pronouncing it dead and overwhelmingly rejected by facts (Kern and Hahn 2018) while others see it as very much alive and kicking (Nei *et al.* 2010; Jensen *et al.* 2019). As a quick glance at major textbooks in population genetics and at the literature would suggest, it seems fair to say that the Neutral Theory is certainly not totally dead. Even if it undoubtedly did lose some of its initial appeal it continues to play a central role in population genetics, a position well summarized by Kreitman (1996) in his spirited essay “The neutral theory is dead. Long live the Neutral Theory”. Shortcomings of the Neutral Theory were already noted in the 1970s and the Neutral Theory has itself evolved. Indeed, its inadequacy to fully explain the data, in particular the constancy of the molecular clock, was already noted in 1973, leading Tomoko Ohta (1973) to propose the Nearly Neutral Theory of molecular evolution. In contrast to the Neutral Theory where most mutations are assumed to be neutral or strongly deleterious, the Nearly Neutral Theory assigns much more prominence to the contribution to standing polymorphism of mutations that are weakly selected and effectively neutral (Ohta 1992; Ohta and Gillespie 1996). Weakly selected mutations can be slightly deleterious or slightly beneficial, but as noted by Kreitman (1996) the best developed of the weak selection models primarily considers slightly deleterious mutations and was therefore christened by him “the slightly deleterious model”. This is the model that we will be testing in most of the present paper.

Like the Neutral Theory, however, the Nearly Neutral Theory still assumes that “only a minute fraction of DNA changes in evolution are adaptive in nature” (Kimura 1983). Under this view, polymorphism is thought to be mostly unaffected by positive selection, except around the few recently selected beneficial alleles (selective sweeps). This was already at variance with the view put forward by Gillespie (e.g. Gillespie 2004) that assigned a greater role to linked positive selection in shaping polymorphism (see also Corbett-Detig *et al.* 2015) and is in even stronger contrast with the claim by Kern and Hahn (2018) that “natural selection has played the predominant role in shaping within- and between-species genetic variation” and that “the ubiquity of adaptive variation both within and between species” leads to the rejection of the universality of the Neutral Theory. In a far more nuanced assessment of the Neutral Theory and its contribution, Jensen *et al.* (2018) argued that the effects of linked selection could readily be incorporated in the Nearly Neutral framework. The heart of the dispute, either today or in the early days of the Nearly Neutral Theory, is about the degree to which each category of mutations contributed directly and indirectly to genetic variation within- and between-species.

A core prediction of the Nearly Neutral Theory is that the fraction of mutations affected by selection depends on N_e_ (Ohta 1973). N_e_ can vary among species but also within a genome because of linked selection (reviewed in Ellegren and Galtier 2016). The effect of selection against deleterious mutations on linked neutral variants – background selection (Charlesworth *et al.* 1993) – is often modeled by a simple re-scaling of N_e_ but except in specific situations effects of linked selection are more complex and there is not a single re-scaling (Barton 1995; Zeng 2013; Comeron 2017; Cvijovic et al. 2018; Torres et al, 2019). In the case of beneficial mutations, for instance, the interference depends both on the beneficial effect of the sweeping mutation and on selection acting at linked sites (Barton 1995; Weissman and Barton 2012).

Evidence that linked positive selection and not only direct selection on slightly deleterious and beneficial mutations contributed to the relationship between the fraction of mutations affected by selection and N_e_ has recently been obtained by Castellano *et al.* (2018). Using two *Drosophila melanogaster* genome re-sequencing datasets, Castellano *et al.* (2018) tested a prediction of the slightly deleterious model first obtained by Kimura (1979) and then extended by Welch *et al.* (2008). Welch *et al.* (2008) showed that if one considers only deleterious mutations, the logarithm of the ratio of nucleotide diversity at non-synonymous and synonymous amino acid changes is linearly related to the logarithm of the effective population size and that the slope of this log-log regression line is equal to the shape parameter of the Distribution of Fitness Effects (DFE), *β*, if the DFE of deleterious mutations is modeled by a Gamma distribution:

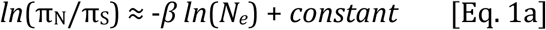

where π_N_ is the nucleotide diversity at non-synonymous sites and π_S_ is the nucleotide diversity at synonymous sites.

Or, rewriting this expectation by using π_S_ as a proxy for N_e_:

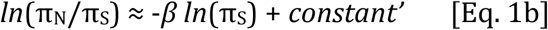

The second equation holds only if variation in π_S_ solely depends on variation in *N_e_*, and that there is no correlation between the mutation rate and N_e_. It should also be pointed out that the DFE used here only considers deleterious mutations, as estimated for instance by DFE-alpha (Eyre-Walker and Keightley 2009). A direct test of this prediction using among-species comparison can be problematic if mutation rates cannot be controlled for. To circumvent this problem, Castellano *et al.* (2018) used within genome variation in *N_e_*, under the reasonable assumption that variation in mutation rates are negligible compared to variation in *N_e_* across a genome. They found (see also James et al. 2017) that the slope was significantly steeper than expected under a simple scaling of N_e_ and simulations indicated that linked positive selection, but not background selection, could explain this discrepancy. The effect of linked selection on the relationship between π_N_/π_S_ and π_S_ is twofold. First it increases stochasticity in allele frequencies, or, in other words, decreases the local effective population size. Second, linked selection leads to non-equilibrium dynamics. Genetic diversity will recover faster for deleterious than neutral mutations, altering the relationship between π_N_/π_S_ and π_S_ (Brandvain and Wright, 2016; Do et al. 2015; Gordo and Dionisio 2005; Vigué and Eyre-Walker 2019). More precisely, the more a region is affected by selective sweeps, the lower π_S_ is and the higher π_N_/π_S_ is compared to the equilibrium expectation: this effect makes the slope steeper compared to the equilibrium expectation.

In the present paper, we first confirmed the observed pattern on the set of 59 species used in Chen *et al.* (2017). We then used 11 high quality genomic datasets for which an outgroup is available to test whether the results obtained by Castellano *et al.* (2018) hold more generally and, in particular, in species with much smaller effective sizes than *D. melanogaster*, and with different levels of linkage disequilibrium. While we adopted the same general approach than Castellano *et al.* (2018), our analysis differed from theirs in one important respect. In their study, Castellano *et al.* (2018) only characterized the DFE of deleterious mutations. We, instead, used a newly developed approach, *polyDFE* (Tataru *et al.* 2017), that also considers positive mutations, which is expected to improve the estimation of the shape of the DFE of deleterious mutations and to disentangle the direct effects of both positive and negative selection.

## Material & Methods

### Genomic data and regression of π_N_/π_S_ over π_S_

In a first step we re-analyzed the 59 species from Chen *et al.* (2017), which included 34 animals and 28 plant species. We estimated the DFE using folded site frequency spectra with the same method as in Chen *et al.* (2017) and calculated the slope (regression coefficient of log(π_N_/π_S_) over log(π_S_) as described in the next paragraph. For DFE estimation using folded SFS the model assumes a gamma distribution for deleterious mutations and takes demography (or sampling or any departure from equilibrium) into account by introducing *n-1* nuisance parameters for an SFS of size *n* (the corresponding code was provided in Chen *et al.* (2017)). In later analyses that required unfolded site frequency spectra, we retained 11 species with high quality genomic datasets and with an available outgroup. These eleven species are given in Table 1. They include both animal and plant species with contrasted levels of nucleotide polymorphism and mating systems. For each of the eleven species, we aligned short reads to the genome using BWA-mem (Li and Durbin 2010) and sorted the alignment using SAMtools. PCR duplicates were removed and INDELs were realigned using GATK toolkit (McKenna *et al.* 2010). HaplotypeCaller was used for individual genotype identification and joint SNP calling was performed across all samples using GenotypeGVCFs. Variant and invariant sites were kept only if genotypes of all individuals were successfully identified (Carson *et al.* 2014). We collected Single Nucleotide Polymorphism (SNPs) in all CDS regions and calculated genetic diversity of 4-fold and 0-fold sites as proxies for polymorphism at synonymous (π_S_) and non-synonymous sites (π_N_). Sites were all masked with ‘N’ and excluded from further computation in the following four cases: heterozygous sites in selfing species, sites with more than two variants, variants at sites within five bases of a flanking INDEL, and missing individuals. We applied the same SNP sampling strategy as in James et al. (2017) and Castellano *et al.* (2018) in order to remove potential dependency between estimates of π_N_/π_S_ and π_S_. In brief, we first split all synonymous SNPs into three groups (S1, S2, and S3) using a hypergeometric sampling based on the total number of synonymous sites. To bin genes and reduce the difference in number of SNPs in each bin, we ranked genes according to their Watterson’s estimate of nucleotide diversity (θ_S1_) and grouped these ranked genes into 20 bins each representing approximately 1/20 of the total number of synonymous SNPs. We then used π_S2_ to estimate the π_N_/π_S_ (mean π_N_ divided by mean π_s_ in each bin) ratio and π_S3_ as an independent estimate of the genetic diversity of each bin.

**Table 1.** Species and datasets used in the present study

| Species | Ref. | Outgroup | Ref. | Mating type | AIC | $b$ | $\beta_{full}$ | $\beta_{gamma}$<br>$a$ | $\beta_{max}$ |
| --- | --- | --- | --- | --- | --- | --- | --- | --- | --- |
| *A. thaliana* | ALONSO-BLANCO *et al.* (2016) | *A. lyrata* | (NOVIKOVA *et al.* 2016) | selfing | 231.3, 227.3 | 0.48 | 0.32 | 0.32 | 0.45 |
| *A. lyrata* | (NOVIKOVA *et al.* 2016) | *A. thaliana* | ALONSO-BLANCO *et al.* (2016) | outcrossing | 247.4, 243.4 | 0.50 | 0.35 | 0.34 | 0.36 |
| *C. rubella* | (KOENIG *et al.* 2018) | *C. grandiflora* | (AGREN *et al.* 2014) | selfing | 201.4, 200.3 | 0.43 | 0.39 | 0.26 | 2.86 |
| *C. grandiflora* | (AGREN *et al.* 2014) | *C. rubella* | (KOENIG *et al.* 2018) | outcrossing | **321.9**, 327.8 | 0.52 | 0.30 | 0.27 | 0.36 |
| *S. habrochaites* | AFLITOS *et al.* (2014) | *S. lycopersicon* | AFLITOS *et al.* (2014) | selfing | **141.5**, 148.1 | 0.21 | 0.23 | 0.13 | 3.61 |
| *S. huaylasense* | AFLITOS *et al.* (2014) | *S. lycopersicon* | AFLITOS *et al.* (2014) | outcrossing | **87.1**, 121.5 | 0.54 | 0.31 | 0.15 | 3.89 |
| *S. propinquum* | MACE *et al.* (2013) | *S. bicolor* | MACE *et al.* (2013) | selfing | 163.8, 159.8 | 0.37 | 0.26 | 0.26 | 0.34 |
| *Z. mays* (teosinte) | CHIA *et al.* (2012) | *T. dactyloides* | CHIA *et al.* (2012) | outcrossing | 208.1, 204.1 | 0.29 | 0.19 | 0.18 | 0.45 |
| *P. trichocarpa* | EVANS *et al.* (2014) | *P. nigra* | (FAIVRE-RAMPANT *et al.* 2016) | outcrossing | 318.9, 319.6 | 0.42 | 0.22 | 0.16 | 2.21 |
| *D. melanogaster* | HUANG *et al.* (2014) | *D. simulans* | STANLEY AND KULATHINAL (2016) | outcrossing | **422.7**, 535.5 | 0.70 | 0.41 | 0.33 | 0.51 |
| *H. timareta* | MARTIN *et al.* (2013) | *H. melpomene* | MARTIN *et al.* (2013) | outcrossing | 208.2, 204.2 | 0.44 | 0.21 | 0.21 | 2.78 |

Note: AIC values were estimated by *polyDFE* for models with and without the effects of beneficial mutations, respectively (bold numbers showed significance < 0.05). The same applies to $\beta_{full}$ and $\beta_{gamma}$ as well. $\beta_{max}$ corresponds to the maximum value of those estimated by *polyDFE* for each ranked gene bin.

We calculated the slope of the linear regression (*l*) of the log-transformed value of the π_N_/π_S_ ratio on the log-transformed value of π_S_, using the “lm” function in R (R Core Team 2018). In pilot runs on 59 species (population data of Chen *et al.* (2017)), the estimates of *l* showed extensive variation depending on, among other things, the qualities of genome sequencing, read depth, annotation and SNP calling. Thus, we selected 11 species for which a high-quality genome sequence and an outgroup were available. Individuals were selected from the same genetic background, i.e. admixture or population structure were carefully removed. At least 20 alleles (i.e. 10 individuals for outcrossing species or 20 for selfing species) were retained from a single ancestral cluster defined in Admixture/Structure analysis in the original publication. For the two *Capsella* species, we performed Admixture analysis for both species separately. A series of quality controls for *l* calculation were performed as described in the following. The longest transcript for each gene model was kept only if it contained both start and stop codons (putative full length) and no premature stop codons. SNPs flanking five bases of INDEL were masked to avoid false positive calls. A grid of filtering criteria (see details in Table S2) was also implemented on each species based on sequence similarity against Swiss-Prot database (e-value, bit-score, query coverage) and sequencing quality (sites with low read depth or ambiguous variants). We selected the filtering criteria in order to maximize the adjusted R^2^ in the log-log regression of π_N_/π_S_ on π_S_. By doing so we aimed to reduce the error introduced by annotation and quality difference between model and non-model organisms. Also, to evaluate the variance introduced by random sampling and grouping of SNPs, we performed 1,000-iteration bootstraps to get the bootstrap bias-corrected mean and 95% confidence intervals for *l* calculations.

**Table 2.** Summary table of multiple regression analyses of the effects of π_S_ *S_b_*, Tajima’s D, and ρ_D_ on Δ, the difference between *b* and *β*.

| $\Delta \sim \pi_S + \log_{10}(S_b)$ | *Coef.* | *SE* | *t value* | *p-value* |
| --- | --- | --- | --- | --- |
| Intercept | 0.14 | 0.031 | 4.69 | 0.0016** |
| $\pi_S$ | 7.93 | 2.96 | 2.68 | 0.028* |
| $\log_{10}(S_b)$ | 0.015 | 3.6e-3 | 4.24 | 0.0029** |
| p-value: 0.0008144 | Adjusted R<sup>2</sup>: 0.7888 |  |  |  |
| $\Delta \sim \pi_S + D + \rho_D$ | | | | |
| Intercept | -0.031 | 0.035 | -0.87 | 0.41 |
| Tajima's D | -0.10 | 0.042 | -2.39 | 0.048* |
| $\rho_D$ | 0.0015 | 6.05e-4 | 2.56 | 0.038* |
| $\pi_S$ | 15.80 | 3.39 | 4.65 | 0.0040** |
| p-value: 0.002978 | Adjusted R<sup>2</sup>: 0.708 |  |  |  |

\*\*\*: $p < 0.001$ , \*\*: $0.001 < p < 0.01$ , \*: $0.01 < p < 0.05$ , ·: $0.05 < p < 0.1$

### Estimates of the distributions of fitness effects

The distribution of fitness effects (DFE) for all non-synonymous mutations across the genome was first calculated by considering only deleterious mutations. We first re-used the DFE parameters estimated in 59 animal and plant species in Chen *et al.* (2017) that assumes that only neutral and slightly deleterious mutations contribute to genetic diversity. In brief, in this previous study the DFE was modeled using a gamma distribution with mean *S_d_* and shape parameter *β*. Folded site frequency spectra (SFS) were compared between synonymous and nonsynonymous sites and demography (or any departure from equilibrium) was taken into account by introducing *n–*1 nuisance parameters for an unfolded SFS of size *n*, following the method proposed by Eyre-Walker *et al.* (2006). The possible issues and merits of this approach compared to those based on an explicit (albeit very simplified) demographic model have been discussed previously and the method introduced by Eyre-Walker *et al.* (2006) has proved to be relatively efficient (Eyre-Walker and Keightley 2007; Tataru e*t al.* 2017). The calculations were carried out using an in-house Mathematica script implementing the method of Eyre-Walker *et al.* (2006) provided in supplementary S2 file of Chen *et al.* (2017).

However, for species with large effective population sizes, like *D. melanogaster*, ignoring the effects of beneficial mutations could distort the DFE to a great extent and lead to a wrong estimate of *β*. Therefore, we further estimated the DFE under a full model that takes both deleterious and beneficial mutations into account (Tataru *et al.* 2017) using unfolded SFS for 11 species. Briefly, the model mixes the gamma distribution of deleterious mutations (shape=*β*, mean=*S_d_*) with an exponential distribution of beneficial mutations (mean=*S_b_*), in proportions of (1-*p_b_*) and *p_b_*, respectively. The unfolded SFS was calculated for the 11 retained species, for which a closely related outgroup with similar sequencing quality was available to polarize the SFS. Ancestral state was assigned as the state of the outgroup if the outgroup was monomorphic for one of the two variants, and the derived allele frequency was calculated from this polarization. Otherwise (in the case of missing data, polymorphic site or third allele in the outgroup) the site was masked. The percentage of SNPs that could not be polarized and were masked varied between 0 and 29.3% with a mean of 4.6% and a median value of 0.5% (Table S2).

In addition, since polarization errors could remain, the error rate of the ancestral state assignment (ε_an_) was also taken into account in *PolyDFE*. The “gamma” DFE (that only considers deleterious mutations) and the full DFE were estimated for each species. In both cases a nuisance parameter was also fitted to account for possible mis-assignment errors in SNP ancestral allele estimation (a step required to obtain the unfolded SFS). Note that, although we used outgroups to polarize SFSs, we did not use divergence but only polymorphism to estimate the effect of beneficial mutations. This is at the cost of larger variance in estimates but it avoids the (potentially strong) bias due to ancient variations in *N_e_* that cannot be captured by modeling recent changes in population size (Rousselle et al. 2018). When comparing the estimates of the DFE among several species, the problem arises that the best model is not necessarily the same for all species (the best model can include or not beneficial mutations and include or not polarization errors). Comparisons cannot be fairly done if all species do not share the same model. Alternatively, estimations under an over-parameterized model can lead to large variance and extreme values. To circumvent this problem, we used a model averaging procedure where each parameter of interest (*β*, *S_b_*, *S_d_*, and *p_b_*) is estimated as a weighted mean of estimates obtained under four models: the Gamma DFE and the full DFE models, including polarization errors or not. The weights given to the estimate from model *k* is wk = e^-1/2ΔAIC_k_^ where ΔAIC*_m_* = AIC*_m_* − AIC*_min_* with AIC being the Akaike Information Criterion and AIC*_min_* the minimum AIC among the four models (Posada and Buckley 2004). All calculations were performed using the software *polyDFE* and the associated R script (Tataru *et al.* 2017).

### Expectations under different selection models

Independently of possible indirect effects of selective sweeps, [Eq. 1] only considers deleterious mutations, in line with the initial view of the Nearly Neutral Theory where beneficial mutations negligibly contribute to polymorphism (Ohta 1973). Giving more weight to beneficial mutations slightly modified the relationship between the slope of the linear regression, *l,* and the shape parameter, *β*. For beneficial mutations only, the equivalent of [Eq. 1] is simply (see Appendix):

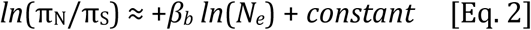

where *β_b_* is the shape of the distribution of beneficial mutations, still assuming a gamma distribution, so *β_b_* would be 1 in the statistical framework we used. Thus, the π_N_/π_S_ ratio increases with *N_e_*, so that considering beneficial mutations the global π_N_/π_S_ decreases more slowly than when only deleterious mutations are taken into account. Thus, with beneficial mutations the slope will always be lower than without. For the majority of species beneficial mutations are rare 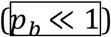 and thus *b* (thereafter we define *b* = −l) is approximately equal to *β*. For those with a relatively high proportion of beneficial mutations, direct positive selection should result in a flattened slope, i.e. a smaller value of *b* than *β*. As we mostly observed the reverse pattern, *b* > *β*, the observed discrepancy cannot be explained by the direct effect of beneficial mutations.

### Trends across the genome and tests for selection

For each of the 20 bins defined above and ranked according to their mean synonymous nucleotide diversity we calculated *β*, *p_b_* and *S_b_* values and a summary statistic of the site frequency spectrum, Tajima’s D (Tajima 1989). Tajima’s D tests for an excess of rare over intermediate variants compared to the frequencies expected under the standard coalescent and was calculated from synonymous sites Demography does affect Tajima’s D and can explain the difference among species. However, a negative Tajima’s D is also expected under recurrent selective sweeps (Jensen *et al.* 2005; Pavlidis and Alachiotis 2017) and should be more negative in genomic regions more strongly affected by linked positive selection. Background selection can also affect Tajima’s D in the same direction but much more weakly (Charlesworth et al. 1995). Independently of the species mean value, we thus expect a strong positive relationship between recombination and Tajima’s D in species where linked positive selection is prominent.

### Forward simulations under selective sweep scenario

The code developed by Castellano et al (2018) which is based on forward simulations using the software SLiM, version 3.2.1 (Haller and Messer 2019) was modified to assess the effect of parameters *p*_b_, *S_b_*, and N on *b* and Tajima’s D. More specifically, a 20-kb genomic region was simulated with a mutation rate of 1×10^-6^ to study the behavior of *b* and Tajima’s D under selective sweep scenarios with varying parameters of *p*_b_, *S_b_*, and N. First, we simulated equal amounts of neutral and deleterious mutations whose fitness effects were drawn from a gamma distribution with a shape parameter 0.4 and a mean *s_d_* of −10. Different percentages of beneficial mutations (*p_b_*= 1%, 0.8%, 0.5%, 0.4%, 0.3%, 0.2%, 0.01%, 0.005% and 0) were drawn randomly from a distribution with a fixed *s_b_* of 1 to simulate loci experiencing selective sweeps at different frequency and we then calculated *b* (Fig. 5 of Castellano et al (2018)) and Tajima’s D. We also investigated the behavior of *b* and Tajima’s D by varying *s_b_* (1, 0.5, 0.1), N (100, 500, 1000) and the recombination rate (Nr=0, 1e-3, 1e-2). Simulated values were averaged across 50 samples, which were taken every 5N generations after an initial burn-in period of 10N generations.

## Results

### b and β are generally similar but the variance is large

One of the most important predictions of the Nearly Neutral Theory is that the proportion of effectively neutral mutations is a function of the effective population size (Kimura and Ohta 1971; Ohta 1972; Ohta 1973; Ohta 1992). In species with large effective population size, selection is efficient and the proportion of effectively neutral mutations is small. Here we used the ratio of genetic diversity at 0-fold over 4-fold degenerate sites (π_N_/π_S_) in protein coding regions as a measure of the proportion of effectively neutral mutations and examined the linearity between log(π_N_/π_S_) and 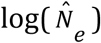 across the genomes of 59 species used in Chen *et al.* (2017). The slope (linear regression coefficient between log(π_N_/π_S_) and 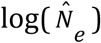 was negative for 51 of the 59 species (*l*<0), although it was significantly different from zero at p=0.05 in less than half of the species (28/59). The value of *l* varied from −0.424 (*D. melanogaster*) to 0.22 (*Callithrix jacchus*) (Table S1). Since balancing selection can lead to both high π_S_ and π_N_/π_S_, it can generate an increase in π_N_/π_S_ for high-π_S_ bins. We thus removed the five bins with the highest diversity and recalculated *l* values for all species. This reduced the *l* values of 36 species and led to negative *l* values in 55 species.

We further examined the DFE for mutations across the genome in the same datasets. A gamma distribution with two parameters, mean (*S_d_*) and shape (*β*), was used to describe the distribution of deleterious mutations under purifying selection. Importantly, the contribution of beneficial mutations, even those under weak selection that are potentially behaving neutrally, is ignored in this case. Estimates of the shape parameter, *β*, varied from 0.01 (*C. jacchus*) to 0.347 (*D. melanogaster*) but were only weakly correlated with effective population size (Table S1).

Considering only deleterious mutations and assuming a simple scaling of N_e_ variation across the genome, the slightly deleterious model predicts that the value of the slope of the linear regression between log(π_N_/π_S_) and 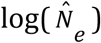, *b* (recall that *b* = −*l*), is equal to *β* (Welch *et al.* 2008). The discrepancy between the two might indicate a departure from this model, and Castellano *et al.* (2018) suggested that in *D. melanogaster,* where the observed slope was steeper than expected, the departure was caused by linked positive selection across the genome. We observed a general consistency between *β* and *b* as estimators of effective neutrality (linear coef. = 1.04, intercept=0.007, p-value<2e-16, adjusted R^2^=0.35, Fig. 1A). The difference (Δ=*b*−*β*) was small in 40 species and varied from −0.1 to 0.1 (Fig. 1B). In 36 species (61%) *b* values were larger than *β* and in 23 species (39%) *β* was larger than *b*. However, the variation in Δ was not explained by π_S_ or N_e_ as the adjusted R^2^ was only 0.06. Removing the five bins with the highest diversity, the correlation between *β* and *b* was still significant (coef. 0.89, p-value=2.14e-6). The median value of Δ increased from 0.0085 to 0.045 but there was still no correlation between Δ and 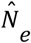.

**Fig. 1.**
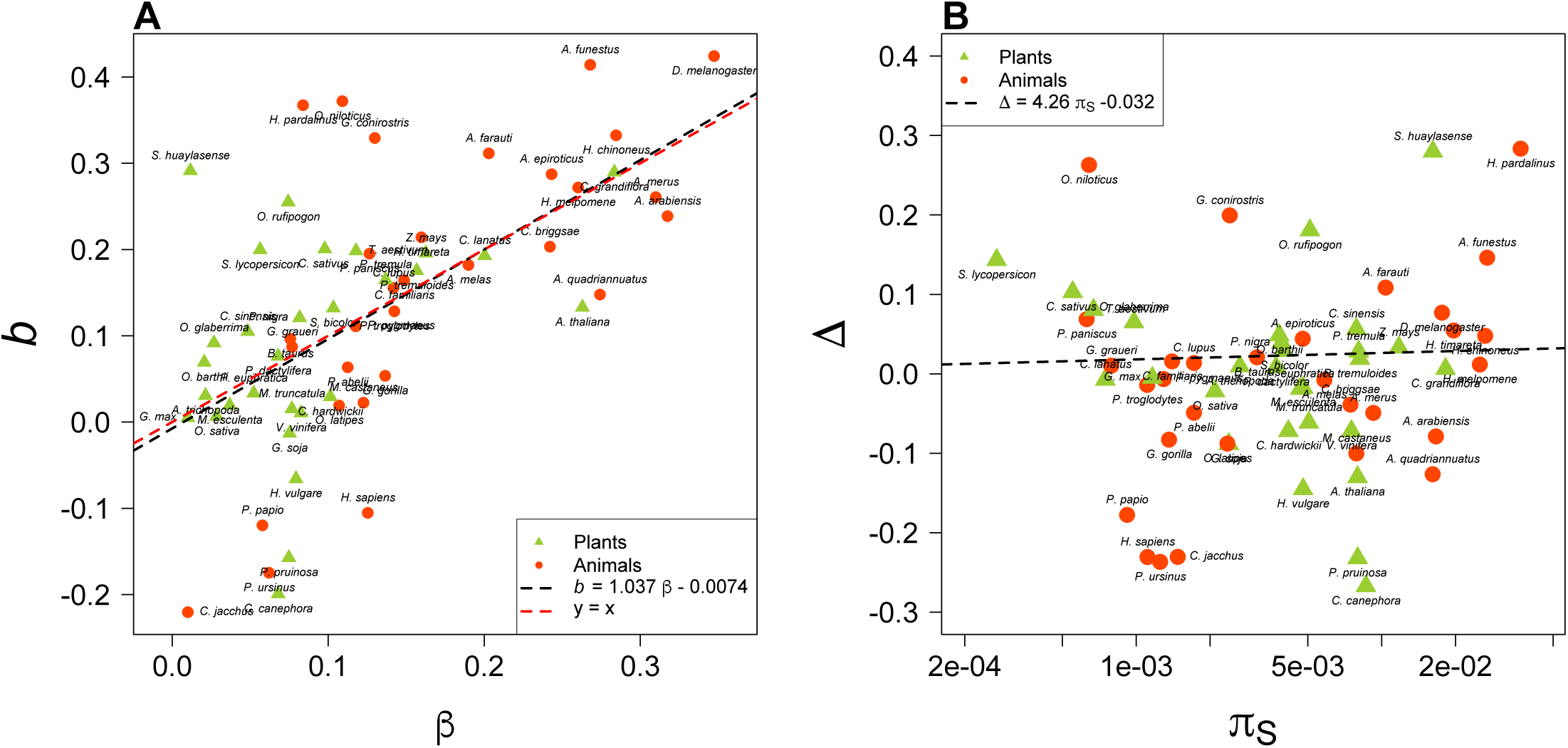
(A) The correlation between *b* and the shape parameter of the DFE, *β,* from the 59 species in Chen *et al.* (2017). The observed slope of the regression of log(π_N_/ π_S_) over log(π_S_), *l*=-*b*. (B) The distribution of Δ (=*b*-*β*) against genetic diversity at synonymous sites. *β* values were estimated from DFE models with only deleterious mutations considered (the gamma distribution).

### The effects of quality control and full DFE model

The variation in Δ may come from two sources. First, it can be due to the estimation quality of *b* and *β*. Tests have shown that quality control on sequencing and SNP-calling can have a dramatic influence on *b* calculations and ignoring beneficial mutations in DFE model could also distort the estimates of *β* (Tataru *et al.* 2017). Second, the variation in Δ can be caused by departures from the assumptions underlying the simple version of the Nearly Neutral Theory, for instance a larger role of direct or linked positive selection than assumed by the theory.

To assess the relative importance of these two sources we selected 11 species with genomic data of high quality and performed a series of stringent quality controls (see details in M&M) before re-estimating *b*. This improved the goodness of fit for the log linear regression between π_N_/π_S_ and π_S_ across the genome and *b* estimates were significantly different from zero for all 11 species (Table 1 and Fig. 2, see also details in Table S2 and Fig. S1). For estimating *β*, we used closely related species to polarize the SFS and applied both the gamma DFE model and the full DFE model implemented in *polyDFE*, which considers both deleterious and beneficial mutations. Instead of choosing the best DFE model, an average value weighted by the different models’ AIC scores was calculated for each parameter (Tataru and Bataillon 2019).

**Fig. 2.**
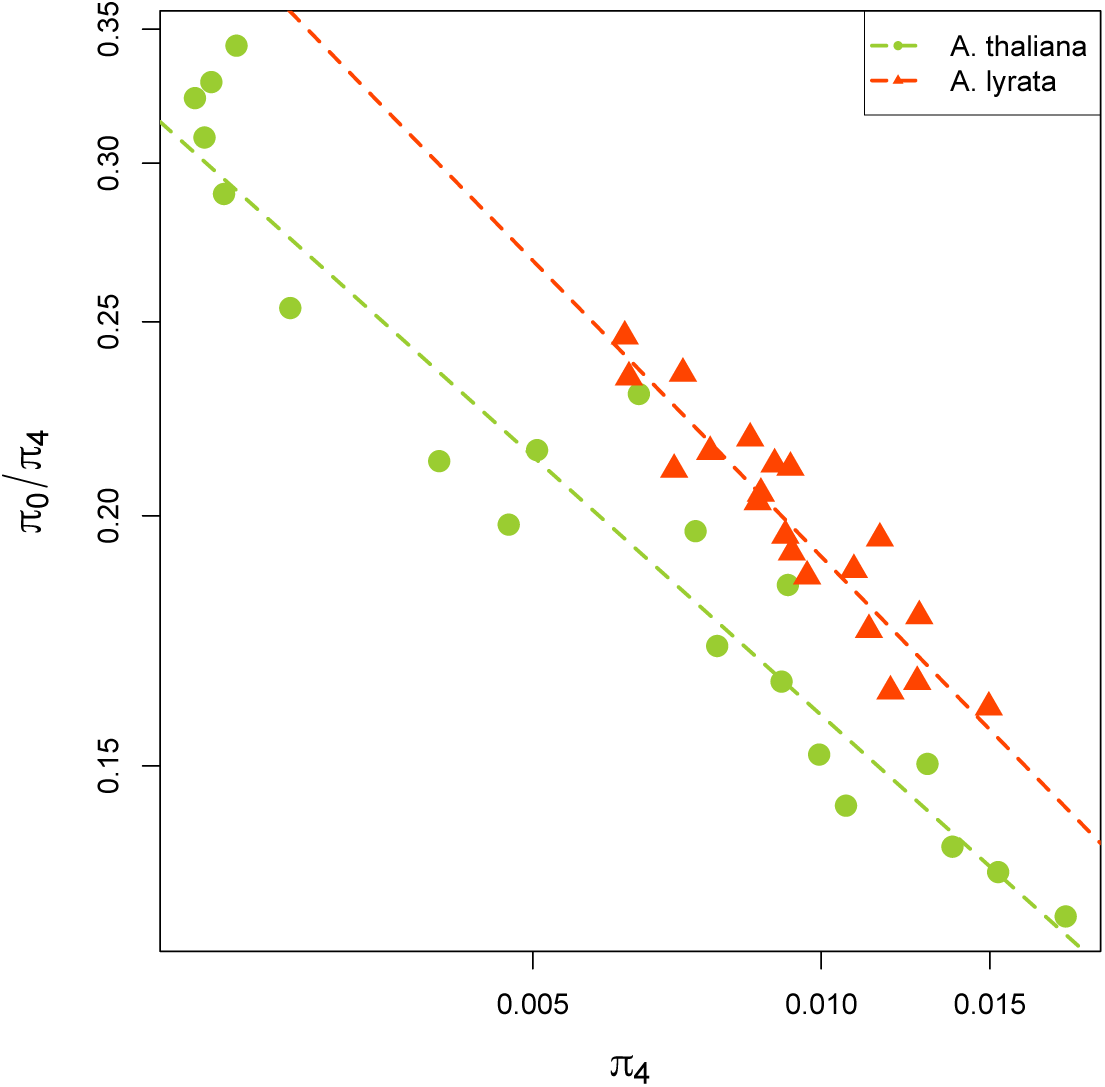
The regression of log(π_N_/ π_S_) over log(π_S_) for self-fertilizing *Arabidopsis thaliana* (dots) and its outcrossing relative *A. lyrata* (triangles).

In this case we observed a better correlation between *b* and *β* (rho = 0.727, p-value=0.011) than when we considered the 59 species and used only a gamma DFE. In addition, considering beneficial mutations slightly increases *β* estimates, making them closer to *b*. However, the linear coefficient between *b* and *β* (1.26) is significantly higher than one and the variation of Δ remains large (−0.026 ∼ 0.289) suggesting that some additional factors may lie behind the remaining variation.

### The roles of effective population size and positive selection

We then tested if the variation in Δ, where Δ=*b*−*β*, could simply reflect differences in effective population size (N_e_) among species. Estimates of *N_e_* were obtained by rescaling π_S_ using estimates of the mutation rate (*μ*) from the literature (see Table S3 for the sources of the *μ* estimates). When Δ is regressed against 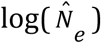, 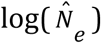 explained up to 49% of the variance in Δ (p-value=0.014). Considering the uncertainty in *μ*, we also regressed Δ on log(π_S_), and obtained similar results (R^2^=0.41, p-value=0.019, Fig. 3).

**Fig. 3.**
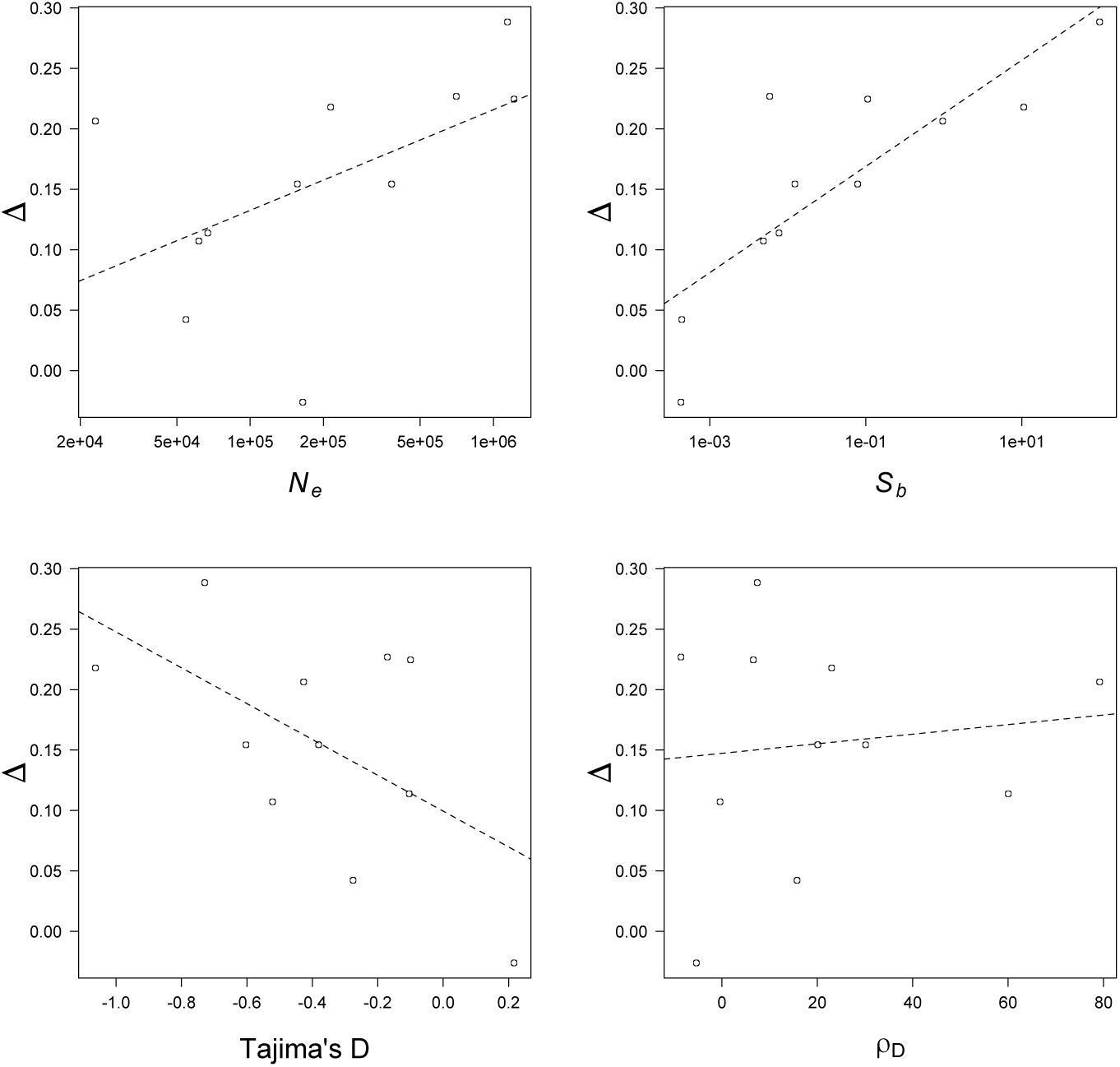
The relationship between Δ (=*b*-*β*) and effective population size, N_e_, selective strength, *S_b_*, Tajima’s D and the trend of D across bins ρ_D_ for 11 selected species. Dotted lines showed the linear regression line. *β* and *S_b_* values were estimated from full DFE models with both deleterious and beneficial mutations considered (full DFE model with both gamma and exponential distributions).

**Table 3.**
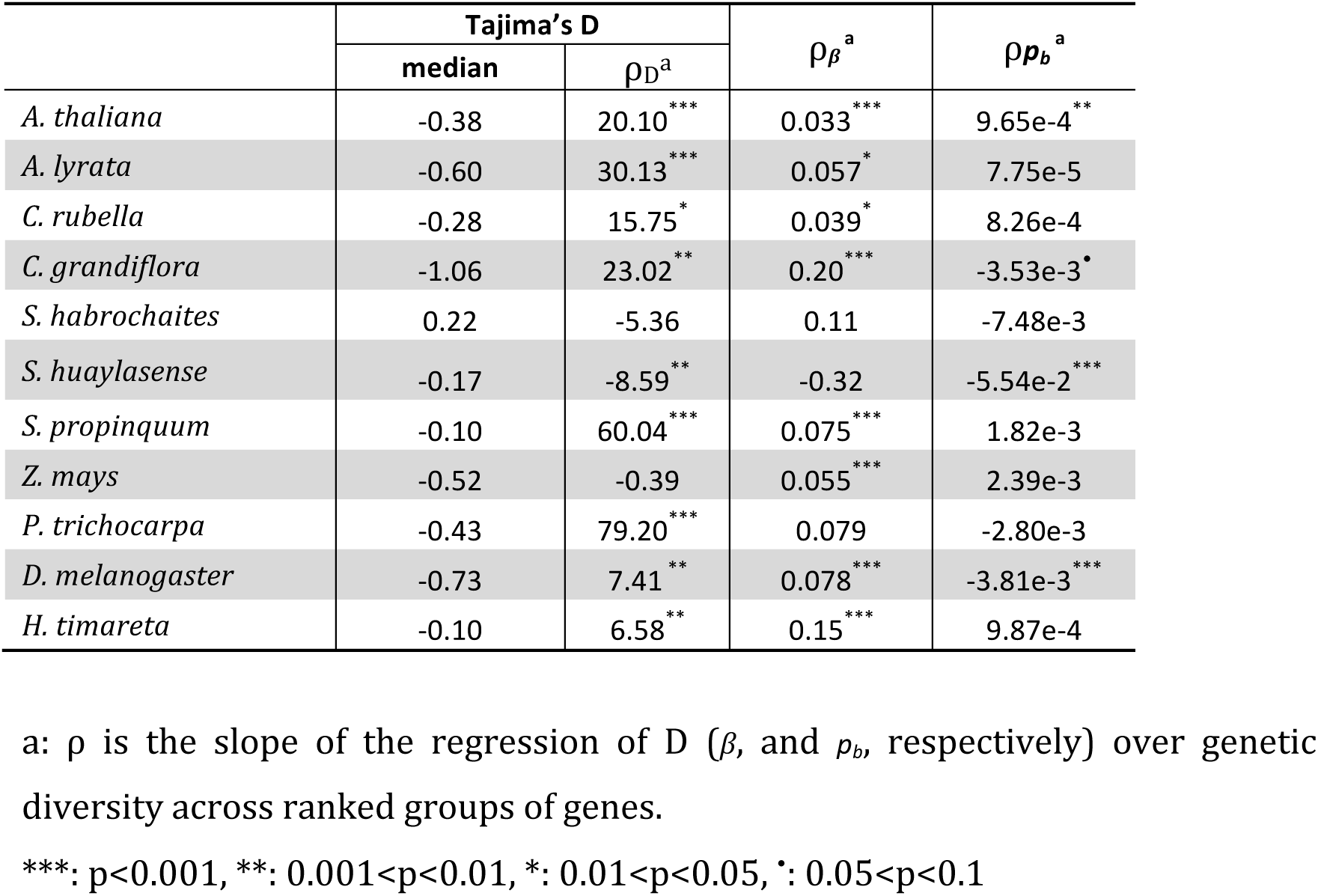
Changes of summary statistics and DFE parameters across 20 rank gene groups.

Furthermore, we tested whether species with potentially more selective sweeps show higher Δ, as predicted by Castellano *et al.* (2018). An explicit model of selective sweeps is difficult to fit given the uncertainty about beneficial mutations parameters and would require additional information, especially on the recombination map of the different species. Alternatively, we qualitatively reasoned that, in addition to be more frequent when the effective population is large, the number of selective sweeps should increase with both the proportion (*p_b_*) and the mean strength of beneficial mutations (*S_b_*). Log(*S_b_*) had a significant and positive effect on Δ (p-value=0.0018, Fig. 3) and explained 64.3% of the variance in Δ but the effect of *p_b_* was not significant (p-value=0.29). When considered together, the effects of both log(*S_b_*) and log(π_S_) (or 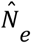) in the joint model explained up to 78% of the variance in Δ (p-value=0.0068 and 0.059, respectively, Table 2). However, no significant effect of *p_b_* could be detected either in the single regression model (p-value=0.29) or joint model with other variables (p-value=0.15). The rate of adaptive evolution relative to the neutral mutation rate, ω_a_ (Galtier 2016) combines the proportion (*p_b_*) and the mean strength of beneficial mutations (*S_b_*) according to ω_a_ = *p_S_* × *S_b_* / (1 – exp(-*S_b_*)). However, as for *p_b_* the effect of ω_a_ on Δ was not significant (p-value=0.17) although the relationship is positive as expected.

### Trends across the genome and tests for selection

Variation of DFE parameters across bins could also explain the difference between *β* and *b* since the underlying assumption is that *β* is constant across bins. We thus calculated *β* for all 20 bins for the 11 species. Seven species had *β* values increasing weakly with genetic diversity (p-value<0.05, mean regression coefficient 0.056) while *C. grandiflora* and *H. timareta* had a much faster increase (regression coefficient =0.2 and 0.15, respectively, Table 3). In five species, the slope was steeper than the maximum *β* value, similar to what was obtained by Castellano et al. (2018) in *Drosophila*. However, the slope was shallower than the maximum *β* value in the six remaining species and in five of them the maximum *β* value was larger than 1 (Table 1). We also compared *p_b_* and *S_b_* values across bins. In *A. thaliana p_b_* increased slowly with diversity whereas in *C. grandiflora*, *S. huaylasense*, and *D. melanogaster p_b_* decreased significantly (p-value<0.05). In all 11 species, *S_b_* did not show any significant trend across bins. To more formally test for the significance of these variations, we also divided the genomes into five bins (to get enough power per bin) and tested the invariance of the DFE across bins using likelihood ratio tests as implemented in *polyDFE*. For all species, a model with independent DFE parameters for each bin is significantly better than a model with shared parameters across bins (see Table S4).

For all 11 selected species we also calculated Tajima’s D (Tajima 1989), thereafter simply called D, in each bin to test for departure from neutrality across the genome. Mean values of D were slightly negative across bins for most species except *S. habrochaites*. For nine of the eleven species, D values increased significantly with genetic diversity (Table 3). Interestingly, we found a negative and strong correlation of Tajima’s D with log(*S_b_*) for all 11 species (p-value=0.0086, Pearson’s correlation coef. =-0.74) but not with any other DFE parameters. This is in agreement with the expectation that selective sweeps decrease D. Background selection could also decrease D albeit to a lower extent. We further tested the trends of positive and negative selection by calculating the proportions of deleterious or beneficial mutations over all bins with selective strength <-10 and >10, respectively. However, no significant trends were identified for either type of direct selection.

We also tested whether alternative measures of the possible occurrence of selective sweeps could explain a larger part of the variation in Δ. We used both the mean Tajima’s D and the among-genome regression coefficient of the relationship between D and π_S_ (ρ_D_) as predictors. More negative D and stronger positive regression coefficient between D and π_S_ can be viewed as signature of stronger hitchhiking effects. So we would expect to see a negative effect of D and a positive effect of ρ_D_ on the variation in Δ. In combination with π_S_ (or 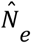), both D and ρ_D_ indeed explained a significant part of the variation in Δ (adjusted R^2^=0.76, Table 2).

### Simulations

Castellano et al. (2018) used forward simulations to assess the extent to which selective sweeps made the slope the relationship between log(π_N_/π_S_) and 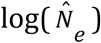 steeper and thereby could explain the discrepancy between the slope and the shape parameter of the DFE, β. They tested varying proportions of adaptive mutations (their Fig. 5). We extended their investigation to test the effect of selective strength (*s_b_*) on *b* with a fixed *β* (0.4) and how selective strength (*s_b_*) also affected estimates of Tajima’s D. Without recombination (Nr=0), Fig. 4 shows that when *s_b_* increased from 0.1 to 1, *b* increased from 0.46 to 0.72 (Δ=0.06 to 0.32). As expected mean Tajima’s D decreased from −0.36 to −0.77 as *s_b_* increased and ρ_D_ between D and π_S_ increased (see also Table 4). We also increased N from 100 to 500, and to 1000, and fixed the mean selective strength at either *S_b_* = 10 or *S_d_* = −1000. With these parameters, the strength of selection was not affected by N but the number of sweeps increased with N due to the higher input of (beneficial) mutations. In this case Δ increased from 0.06 to 0.41 as N increased and Tajima’s D again decreased (Table 4 and Fig. 5). With recombination (Nr=1e-3 and Nr=1e-2), we noticed similar trends of *b*, D, and ρ_D_ when *s_b_* or N are large enough to recover the significance of the linearity between log(π_N_/π_S_) and log(π_S_) (Fig. S2 and S3).

**Table 4.**
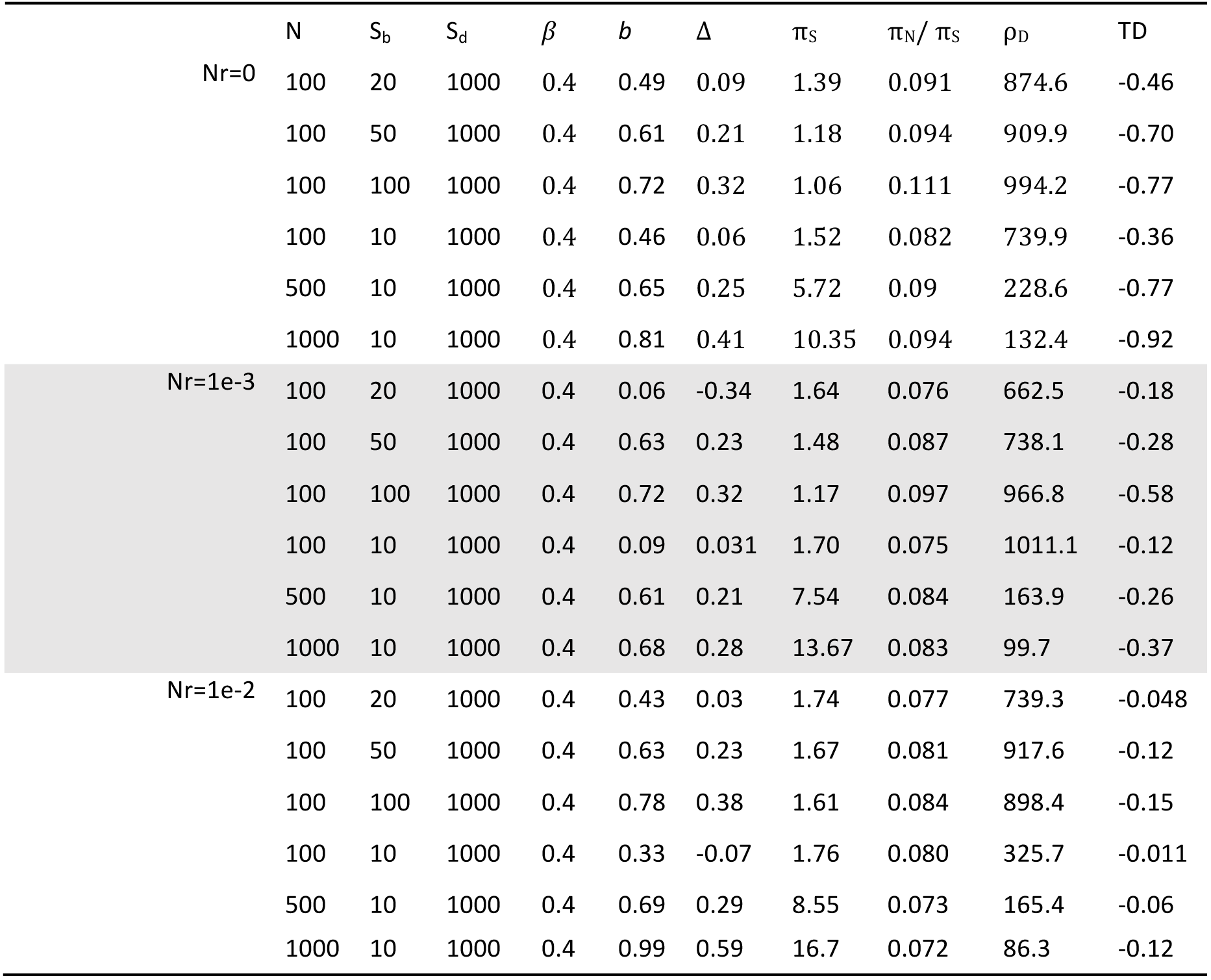
Results of forward simulations showing the effect of linked positive selection on *b*, Δ and summary statistics of the site frequency spectrum for different values of the mean selective value of beneficial mutations, S_b_ and the population size, N. ρ_D_ is the correlation between π_S_ and Tajima’s D.

**Fig. 4.**
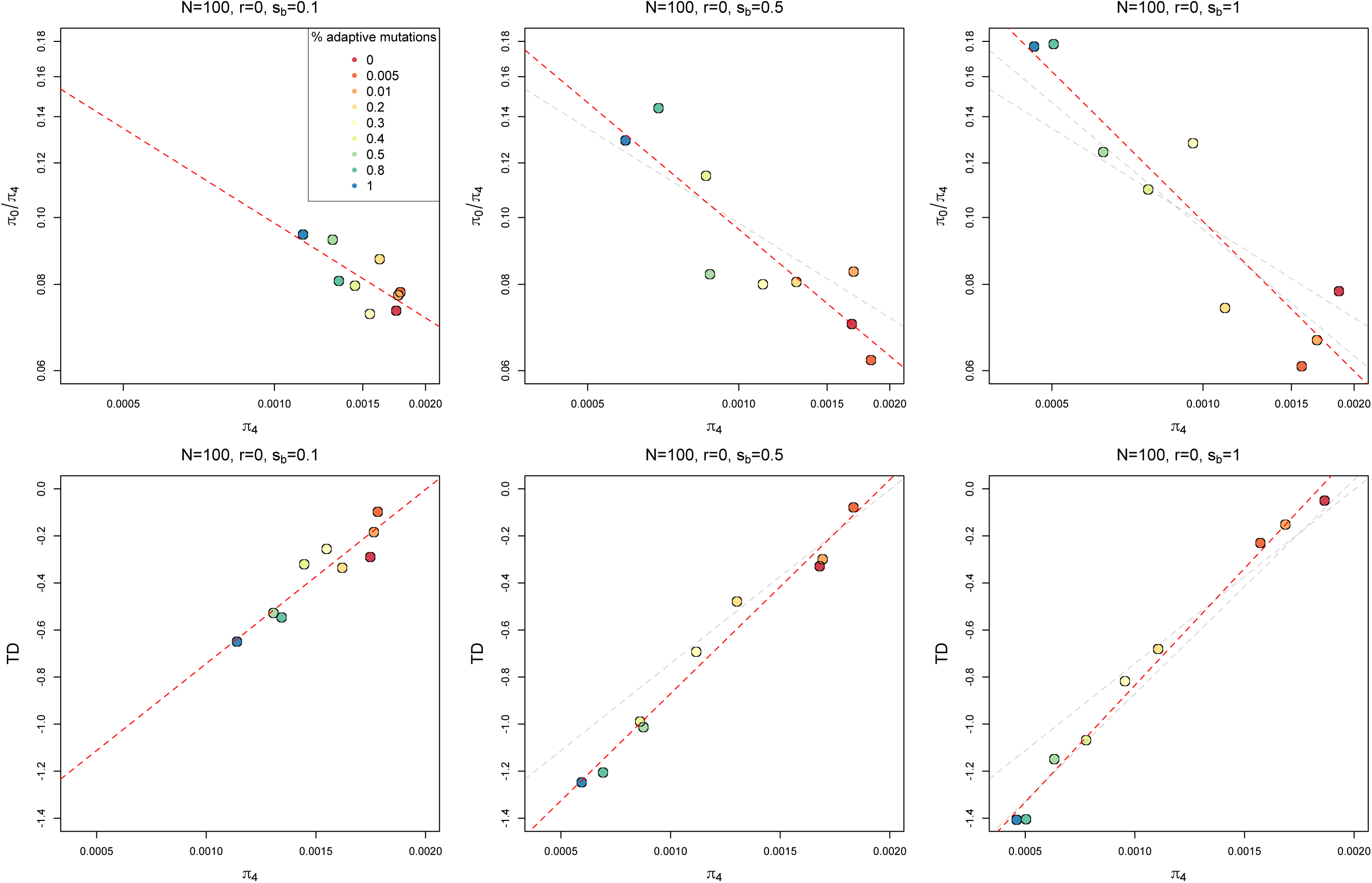
Effect of linked positive selection on the relationship between log(π_N_/π_S_) and log(N_e_) and Tajima’s D. Upper row: The linear regression coefficient (*b*) between log(π_N_/π_S_) and log(N_e_) increases with increasing positive selective strength (from left to right). The red lines are the regression lines for each case. To facilitate comparisons among figures, and illustrate how the slope gets steeper as *s_b_* increases the regression lines corresponding to s_b_=0.1 and/or s_b_=0.5 values are reported with gray lines. Lower row: The red lines for Tajima’s D panels indicate the mean values.

**Fig. 5.**
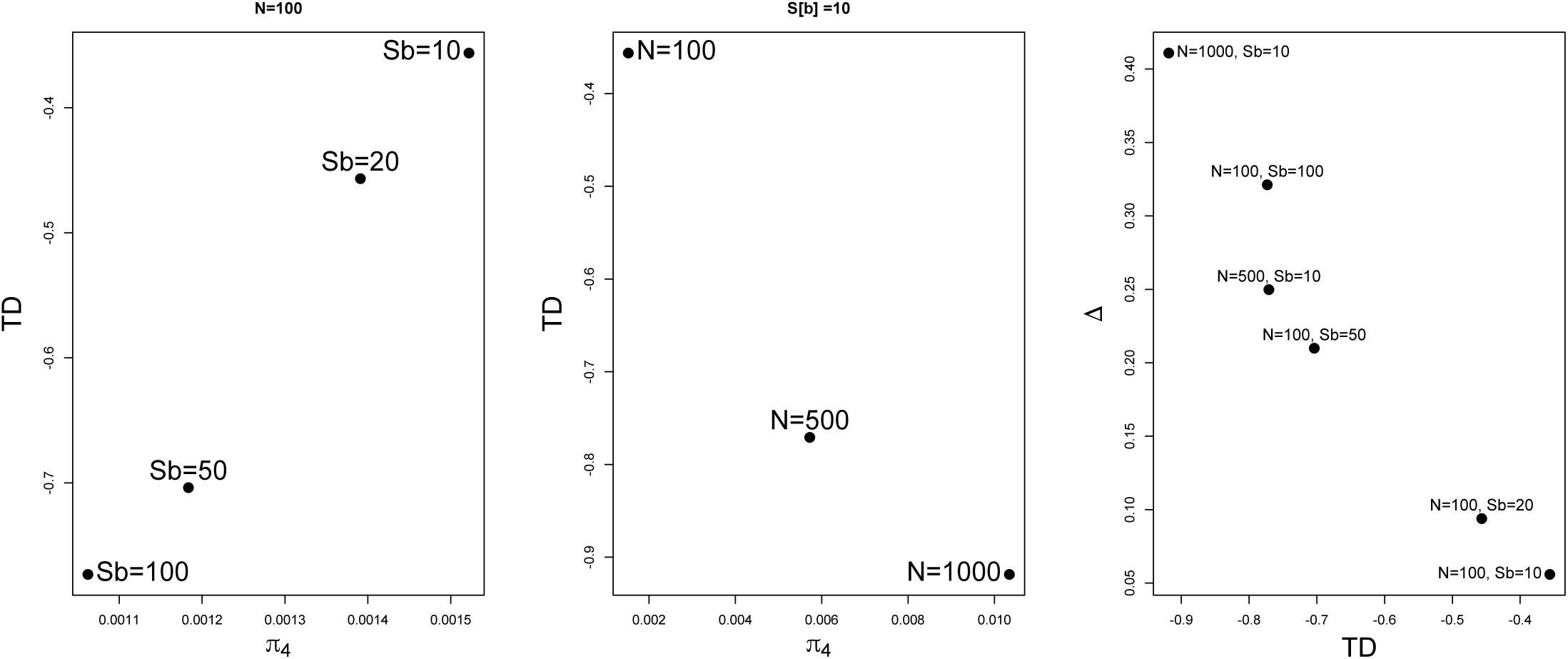
The correlation between Tajima’s D and π_S_ depending on S_b_ (left panel) and N (middle panel); Correlation between Δ and Tajima’s D (right panel). In all three cases the results were obtained with forward simulations in Slim assuming no recombination.

## Discussion

The aim of the present study was to test quantitatively one of the predictions of the Nearly Neutral Theory of molecular evolution or, more precisely, the slightly deleterious model. More specifically, we used full genome datasets to test whether the proportion of effectively neutral mutations varies with local variation in N_e_ across the genome and decreases linearly with increasing N_e_ and whether the slope is equal to the shape parameter of the DFE. The negative log linear relationship between π_N_/π_S_ and N_e_ observed in previous studies (Gossmann et al. 2011; Murray et al. 2017; Castellano et al. 2018; Vigué and Eyre-Walker 2019) was also observed in the present study, although the slope was not always significantly negative and, when negative, could differ significantly from the shape parameter of the DFE and be much steeper. The latter was especially true in species with large effective population size and the difference was correlated to the estimated mean strength of selection acting on beneficial mutations. In the case of species with large effective population size neglecting linked positive selection could therefore lead to a significant quantitative discrepancy between predictions and observations. On the other hand, the slightly deleterious model appears as a good approximation when the effective population size is small. Below we first consider possible caveats and discuss the implications of the results for the relative importance of purifying and adaptive selection in shaping the genetic diversity of species.

The discrepancy between the slope of the log linear relationship between π_N_/π_S_ and N_e_ and *β* could simply be due to difficulties in estimating them precisely. In general, estimates of the DFE shape parameter, *β,* were rather stable compared to estimates of the slope of the regression of log(π_N_/π_S_) over log(π_S_), *b*, with the variance of the former being half that of the latter independently of quality control and whether the SFS was folded or unfolded. High variation in *b* estimates may explain the fact that a significant correlation between π_N_/π_S_ and π_S_ could not be observed for all species, particularly those with low genetic diversity (e.g. great apes). Therefore, a stringent quality control for read alignment and SNP calling is necessary, even for *D. melanogaster*, where an improvement of the fit in *l* calculation (linear regression adjusted R^2^=0.79 to 0.95) leads to a dramatic change in the estimate of Δ (from 0.077 to 0.29). Even if a stringent quality control had been implemented, the goodness of fit for the log linear regression leading to the estimation of *b* would differ significantly from species to species. The fit across the *D. melanogaster* and *A. thaliana* genomes was almost perfect (R^2^>0.95) while, at the other extreme, the fit was rather poor in *S. habrochaites* (R^2^=0.38). However, even among species for which the fit is almost perfect (R^2^>0.95) *b* could vary rather dramatically: *D. melanogaster* had a much larger *l* (0.7) than *A. thaliana* (0.48), *C. rubella* (0.43), and *Z. mays* (teosinte, 0.29), whereas *β* only changed marginally for these species. Not all species though showed a significant negative linear relationship between π_N_/π_S_ and 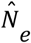 and some even had positive slopes, especially for those of low diversity (e.g. great apes, Fig 2). Therefore, besides purifying selection the slope is also likely to be affected by additional factors. Factors that affect the likelihood to observe a negative relationship between π_N_/π_S_ and 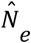 and its relationship with the DFE parameters were thoroughly discussed by Castellano et al. (2018). Below we highlight those that seem particularly relevant when considering a group of species with contrasted levels of diversity as was done here. These factors are the variation in N_e_ estimates along the genome, which itself reflects the joint distribution along the genome of recombination rate and density of selected sites, the DFE, and the variation along the genome of the rate of adaptive evolution (Castellano et al. 2018).

Lack of joint variation in recombination rate and selected sites seems to be an unlikely cause for an absence of negative relationship between π_N_/π_S_ and N_e_ as such a relationship is observed in selfing species where this joint variation is expected to the more limited than in outcrossing ones. A possible source of variance in *β* could be that the single-sided gamma distribution does not describe well the real DFE curves, at least not for all species, particularly when the DFE is not unimodal (Tataru *et al.* 2017). For species like *D. melanogaster*, for instance, there is mounting evidence of adaptive evolution (reviewed in Eyre-Walker 2006, Sella et al. 2009). Therefore, it is necessary to consider the possible contribution of beneficial mutations. The full DFE model provided a much better fit than the gamma DFE that considers only deleterious mutations in *D. melanogaster* (log likelihood= −187.3 versus −245.7, respectively). This was also true of some of the outcrossing plants like *Capsella grandiflora*, and *Solanum huaylasense*. In all three species *β* estimates increased when estimated with the Full DFE instead of the Gamma DFE, sometimes significantly (from 0.33 to 0.41 in *D. melanogaster* (Rwanda) and 0.15 to 0.31 in *S. huaylasense*) and at other times only marginally (0.27 to 0.30 in *C. grandiflora*). Taking beneficial mutations into account when fitting the shape of the DFE can partly reduce the discrepancy between *β* estimates and the slope of the regression. However, it is not sufficient as Δ was positive in 10 over the 11 focal species we studied.

Based on the prediction of the Nearly Neutral Theory with direct positive selection (Equation 2), the proportion of beneficial mutations is the only factor that could alter the relationship between *b* and *β* and should always result in a larger *β* compared to *b*. However, this is usually not the case as, on the contrary, values of *b* larger than *β* have generally been reported (Chen *et al.* 2017; James *et al.* 2017; Castellano *et al.* 2018). In this paper we systematically investigated this relationship across the genomes of multiple species. Two thirds of the 59 species and 10 out of the subset of eleven species that were selected for the high quality of their genome, had larger *b* than *β* values. Hence direct positive selection is not the main cause of the discrepancy.

Investigation of DFE parameter changes across bins may help to identify changes in natural selection. Increasing *β* values over bins could be a signal for stronger positive selection in low diversity regions. Although the maximum *β* value of some species can be larger than *b*, *β* grows slowly for most species and shows hardly any pattern between species. Neither did *p_b_* or *S_b_*. This lack of significant trend in these parameters could simply be due to an increase in variance of their estimates as only one twentieth of the total number of polymorphic sites were used for DFE calculations in each bin. It could also again suggest that direct selection is not the main cause of the discrepancy.

One of the main findings of the present study is that a large proportion of variance in the discrepancy can be explained by the estimated strength of positive selection, which can be regarded as an indication for linked selection, such as selective sweeps or more generally hitchhiking effects. To test for that, we compared changes in Tajima’s D and its among-genome correlation coefficients over bins. As expected we observed a negative effect of D and a positive effect of ρ_D_ on Δ, both suggesting the presence of linked selection, with lower diversity at nearby sites and thus increased discrepancy between *b* and *β*. This is also in agreement with our simulations and those of Castellano et al. (2018) that illustrate that hitchhiking effects can lower the genetic diversity at nearby neutral or nearly neutral positions. These results can be understood because selective sweep effects cannot simply be captured by a rescaling of N_e_. Selective sweeps not only reduce genetic diversity at linked sites but also distort the coalescent genealogy (Fay and Wu 2000; Walsh and Lynch 2018; Campos Parada and Charlesworth 2019), so that we cannot define a single N_e_ in this context (Weissman and Barton 2012). In particular, the scaling is not expected to be the same for neutral or weakly selected polymorphisms. However, as far as we know, there is no quantitative model predicting the value of the slope as a function of DFE, rates of sweep and recombination rates, and such models still need to be developed.

## Conclusions

There are three major conclusions to the present study. First, the Nearly Neutral Theory in its initial form may not explain all aspects of polymorphisms but, almost 50 years after it was first proposed by Tomoko Ohta (Ohta 1973), it still constitutes an excellent starting point for further theoretical developments (Galtier 2016; Walsh and Lynch 2018). Second, considering linked beneficial selection indeed helps to explain more fully polymorphism data, and this is especially true for species with high genetic diversity. This can explain both patterns of synonymous polymorphism (Corbett-Detig et al. 2015) and how selection reduces non-synonymous polymorphism (Castellano et al. 2018, this study). One could have a progressive increase of the effect of selective sweeps as suggested by Walsh and Lynch (2018, chapter 8) with a shift from genetic drift to genetic draft (Gillespie 1999; 2000; 2001). If so, we could have three domains.

For small population sizes, drift would dominate and the nearly neutral theory in its initial form would apply. For intermediate population sizes beneficial mutations would start to play a more important part, and finally for large population sizes, the effect of selective sweeps would dominate and draft would be the main explanation of the observed pattern of diversity. Third, our study once more emphasizes the central importance of the DFE in evolutionary genomics and we will likely see further developments in this area.

## Acknowledgements

We thank Thomas Bataillon and David Castellano for comments on earlier versions of the manuscript. The project was in part supported by grants from the Swedish Research Council and the Swedish Foundation for Strategic Research to ML.

## Data availability

The vcf files used in the present study are available on request.

## Supplementary Information

**Supplementary table**

**Supplementary table legends**

Table S1. The 59 species used to compare the difference between *-l* and *β* assuming a gamma model for DFE. See Chen et al. (2017) for further details.

Table S2. Details of the 11 species used in the current study to compare the difference between *-l* and *β* assuming a full model (gamma + exponential) for the DFE.

Table S3. Mutation rates used for 11 species used in the current study for estimation of N_e_.

Table S4. Test for the invariance of DFE parameter estimates across bins by comparing the log-likelihoods of independent estimates for each bin against those of shared estimates.

## APPENDIX

In a constant population with population size *N_e_*, π_S_ = 4*N_e_µ* and π_N_ is given by (Sawyer and Hartl 1992):

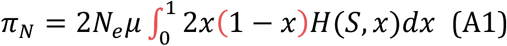

where

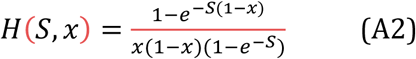

is the mean time a new semidominant mutation of scaled selection coefficient *S* = 4*N_e_s* spends between *x* and *x* + *dx* (Wright 1938). For constant selection *S*, by integrating (A1) and dividing by 4*N_e_µ*, we have:

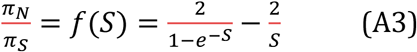

(A3) is valid for both positive and negative fitness effect. If we consider only beneficial mutations with a gamma distribution of effects, with mean *S_b_* and shape 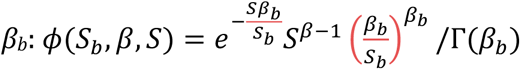, we can use the same approach as Welch et al. (2008) to show that:

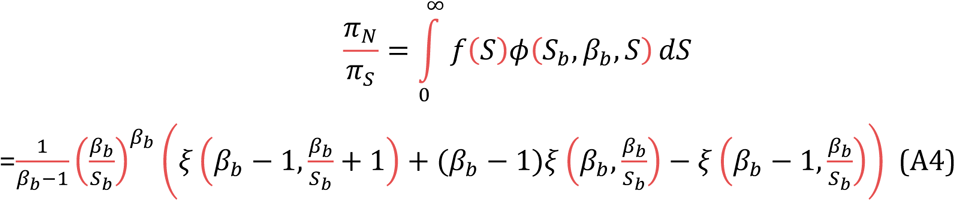

where *ξ*(*x*, *y*) is the Hurwith Zeta function. (A4) can be approximated under the realistic assumption that 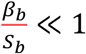 and taking Taylor expansion of (A4) in 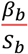 around 0. We thus obtain:

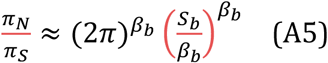

which leads to equation [eq. 2] in the main text.

